# REFLEX, a Novel Immune Profiling Assay, Combining TCR Repertoire and Multiome at Massively Scalable Single-cell Resolution to Catapult Exploration of T-cell Derived Immunity

**DOI:** 10.1101/2025.10.24.684243

**Authors:** Matthew R. Hart, Zachary J. Thomson, Saransh N. Kaul, Saskia Ilcisin, Matthieu Landreau, Tyanna J. Stuckey, Peter J. Wittig, Micheal Keller, Chase D. McCann, Troy R. Torgerson, Peter J. Skene

**Author notes:** These authors contributed equally to the publication (Matthew R. Hart, Zachary J. Thomson).

## Abstract

Single-cell profiling of T cell state with immune repertoire is critical for understanding heterogenous T cell phenotypes and responses to antigen, however, existing technologies struggle to generate this information at sufficient throughput to match biological complexity. We present “REFLEX”, a novel single-cell method enabling highly scalable, cost-efficient, multiomic profiling with paired-chain TCR sequencing. REFLEX utilizes in-situ reverse transcription with integrated sample multiplexing barcodes in a way that merges seamlessly with the commonly used 10x FLEX platform to allow capture of TCR sequences at unprecedented scale and depth. We profile >2 million cells from CMV-peptide-pulsed T cell expansions, capturing TCR sequences and rich multiomic information from 1.4M T cells, identifying many putative novel CMV reactive clonotypes and illustrating the scale and transformative impact on our understanding of T cell mediated adaptive immunity achievable with REFLEX.

## Introduction

T-cell receptor (TCR) sequencing is a cornerstone technology in the study of T cell/antigen interactions. While bulk RNAseq profiles TCR diversity and abundance at the sample level, merging paired α/β chain and cell-state information requires single-cell (sc) resolution^1^. Recent benchmarking studies of scRNAseq platforms have concluded that droplet-based offerings outperform others by most metrics^2,3^. Of these, 10x Genomics 5’ platform is TCRseq compatible but lacks the scalability to address the inherent complexity of the search space with a predicted repertoire of >2.5×10^7^ unique TCRs per person^4,5^.

The 10x FLEX platform enables pre-GEM (Gel-Beads-in-Emulsion) sample barcoding and multiplexing, resolvable hetero-multiplets, and exponentially increased scalability over 10x 5’. Its chemistry allows diverse sample types which can be collected, fixed, stored, and later processed in parallel, saving time/cost and enabling logistically difficult longitudinal or geographically diverse study formats^6^. FLEX measures gene expression (GEX) by quantifying hybridized/ligated probe pairs rather than through capture of reverse transcribed cDNA. Thus, it fails to generate the underlying genetic sequence information required for TCR study. Here, we present REFLEX (REpertoire-FLEX), a method designed to supercharge the exploration of T cell immunity by enabling single-cell capture of TCRα/β sequence on a highly scalable, cost-effective, multi-modal platform (Fig1a).

**Figure 1.**
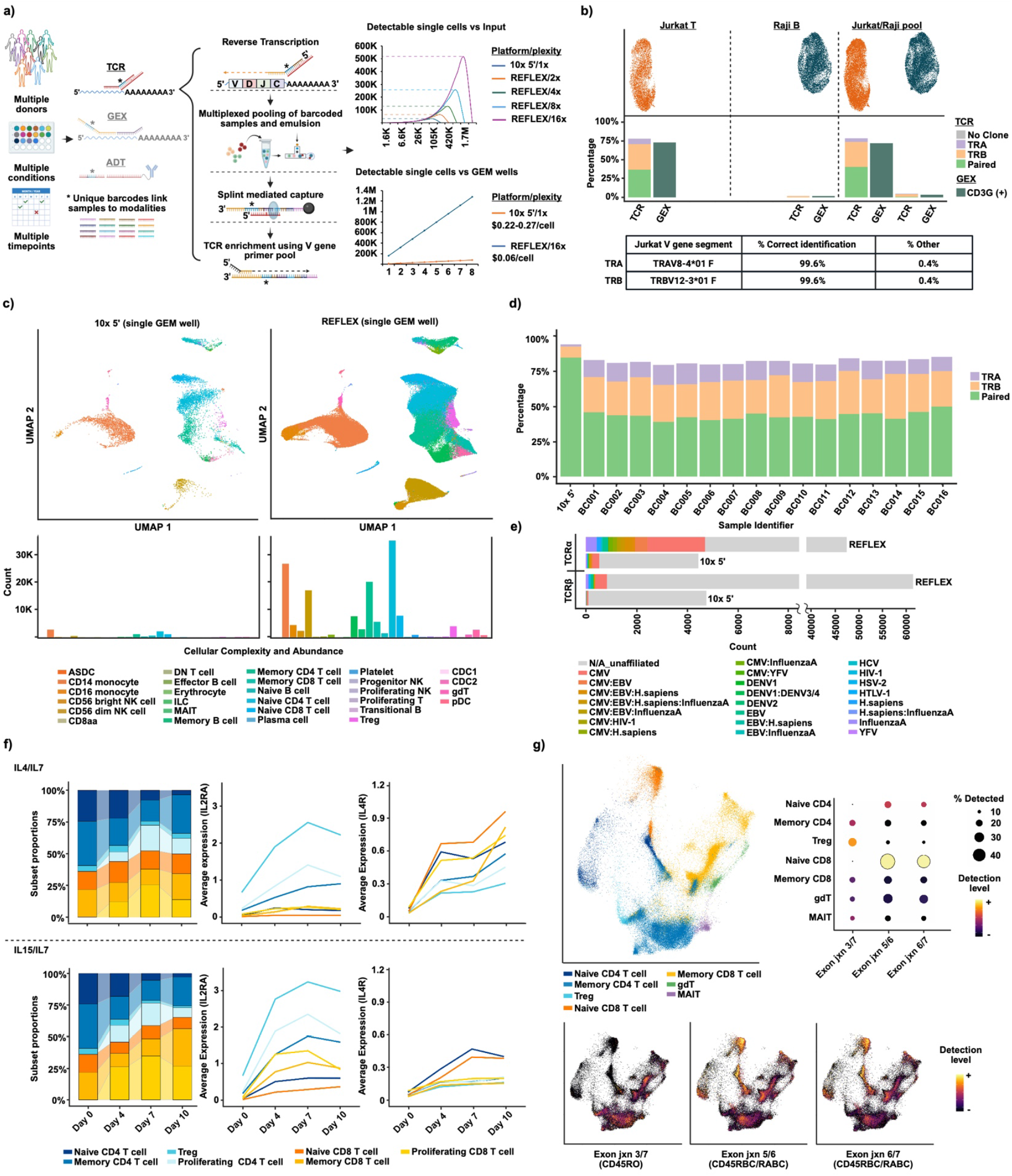
REFLEX enables massively scalable capture of single-cell TCR repertoire and transcriptome on the highly versatile FLEX platform. a) Schematic of REFLEX compatibilities and workflow; performance compared to 10x 5’ assay. Graphs show theoretical capture from “superloading” (upper) and cost/cell with standard loading by GEM well (lower). b) Sensitivity and specificity of α/β/paired capture in pooled and individual T and B cell lines. Table reflects Jurkat TCR V-segments identified. c) PBMC cell typing and abundance in single-well captures by REFLEX vs 10x 5’. d) Paired and single chain capture rates across 16 barcodes (REFLEX) vs 10x 5’. e) Quantified TCRα/β captured by 10X 5’ and REFLEX, showing CDR3 epitope association by VDJdb. f) T-cell subset proportions and selected GEX in response to culture with IL4/7 or IL15/7 and g) UMAPs showing detection of custom CD45 isoform probes from day 7, IL4/7 and IL15/7 combined.

## Results

REFLEX captures TCRα/β utilizing primers targeting invariant 3’ “constant” regions, *TRAC* and *TRBC*. 5’ primer overhangs encode 10x sample barcodes (permitting pre-GEM sample multiplexing) as well as a constant sequence allowing hybridization to pre-annealed DNA splinting oligos (SuppTables1,2). In-situ reverse transcription is performed following overnight probe hybridization, generating single-stranded cDNA spanning the VDJ inclusive, complementarity determining region 3 (CDR3). Capture and ligation of cDNA onto gel bead oligos during in-GEM incubation is mediated by the splinting oligos mentioned above. During pre-amplification, a pool of Variable-segment primers (SuppTable3) provides enrichment of full-length CDR3. Pre-amplified cDNA is then partitioned and further prepared as separate libraries by modality (SuppFigs.1,2). Sequencing/analysis utilizes standard, open-source packages/pipelines (Methods).

REFLEX specificity and accuracy were validated using clonal Jurkat-T and Raji-B cell lines. Within Jurkats, REFLEX detected literature reported V-segments with >99% accuracy^7^(Fig1b). Multiplexed pooling of both produced only trace detection of TCR in B cells which was comparable to that of FLEX probes for T cell marker, *CD3G*, suggesting low-level ambient RNA contamination intrinsic to single-cell technologies (Fig1b). We next evaluated consistency across sample barcodes in the context of human PBMCs. Technical replicates were processed with 16 unique sample barcodes, pooled, and loaded into a single GEM well at the maximum recommended input (256k cells @ 16k/barcode). We observed consistent detection of genes and UMI/cell across barcodes (median gene detection range: 1874-2195, MAD = 63.38, median UMI range: 2963-3701.5, MAD = 149.74, SuppFig.3). Importantly, the superior scaling capabilities of REFLEX provided a >15-fold increase in cells captured over an individual well of 10x 5’ (145,677 vs 9,453 cells^8^), enhanced resolution of PBMC heterogeneity, and superior identification of minor cellular subtypes (Fig1c). TCR performance across barcodes was also highly consistent (mean paired TCR = 43.6%, mean **β** = 70.2%, mean α = 55.6%, Fig1d) with cumulative TCRα/**β** chain detection exceeding 10x 5’ by nearly 10^5^ clones (Fig1e). While several thousand of these clones were associated with known epitope species within online repositories such as VDJdb ^9,10^, most were orphan in nature highlighting sparsity within existing databases and the utility of highly scalable approaches for identifying TCR/antigen pairings.

The adaptability of the REFLEX extends beyond scalability. Fixation compatibility enables parallel processing of longitudinally harvested samples, reducing batch effects and cost. Additionally, custom FLEX probe supplementation to the existing >18,000 gene panel is simple and cheap, facilitating exploration of difficult-to-address spliceoforms or non-polyadenylated transcripts, etc.^11^. To demonstrate these functionalities with REFLEX, we designed probes targeting isoform defining exon/exon junctions of *PTPRC*/*CD45* (SuppFig.4). We cultured PBMCs from 3 donors over 10 days in the presence of IL-4/IL-7 or IL-15/IL-7, shown in-vitro to promote homeostatic proliferation of CD4 or CD8 T-cells, respectively^12,13^. Aliquots were taken at days 4/7/10 and fixed/permeabilized/frozen at -80C. Samples were processed with REFLEX, generating TCR, GEX (SuppFig5), and *CD45* isoform information (Day 7 only). In line with published literature, we observed enrichment of CD8 memory and proliferating T cells in IL-15/IL-7 cultures relative to IL-4/IL-7 (Fig1f, Methods) along with elevated cytokine dependent gene expression (*IL2RA* vs *IL4R*). As anticipated, custom probes spanning exons 3-7 (*CD45RO*) or 5-6 and/or 6-7 (*CD45RBC*/RABC) were enriched in memory and naïve subsets, respectively^13^ (Fig1g). These results demonstrate compatibility of REFLEX TCR with fixation/storage and GEX/custom FLEX probe capabilities.

REFLEX was developed to bridge gaps in areas of research or medicine where throughput requirements and current technologies are at odds. For example, adoptive transfer of expanded/enriched virus specific T cells (VSTs) has shown efficacy in preventing refractory viral infection post hematopoietic stem-cell transplant^14^ (Fig2a). Unfortunately, comprehensive understanding of VST variability from donor heterogeneity or production method is lacking given standard approaches and scRNAseq +/-TCR is currently impractical due to throughput limitations^15^. To evaluate REFLEX scalability and fixation advantages in this context, we generated VSTs from CMV seropositive donors using a clinically tested approach^16^. PBMCs, pulsed with CMV derived peptide pools, were cultured with IL-4/IL7 or IL-15/IL-7 over 10 days (Methods). CMV reactive T-cell expansion was initially confirmed using fluorescent HLA matched dextramers (Methods, SuppFig6). Aliquots, harvested throughout culture, were fixed/permeabilized/stored at -80C for longitudinal profiling. Samples were processed in parallel, by donor, in three multiplexed REFLEX runs, each with 2 GEM-well replicates (∼512,000 cells/donor). REFLEX detected an average of 64.1% paired TCR clones, with 74.7/75.5%, α/β, across the 3 donors with optimal detection of paired or single clones at later timepoints (SuppFig7).

**Figure 2.**
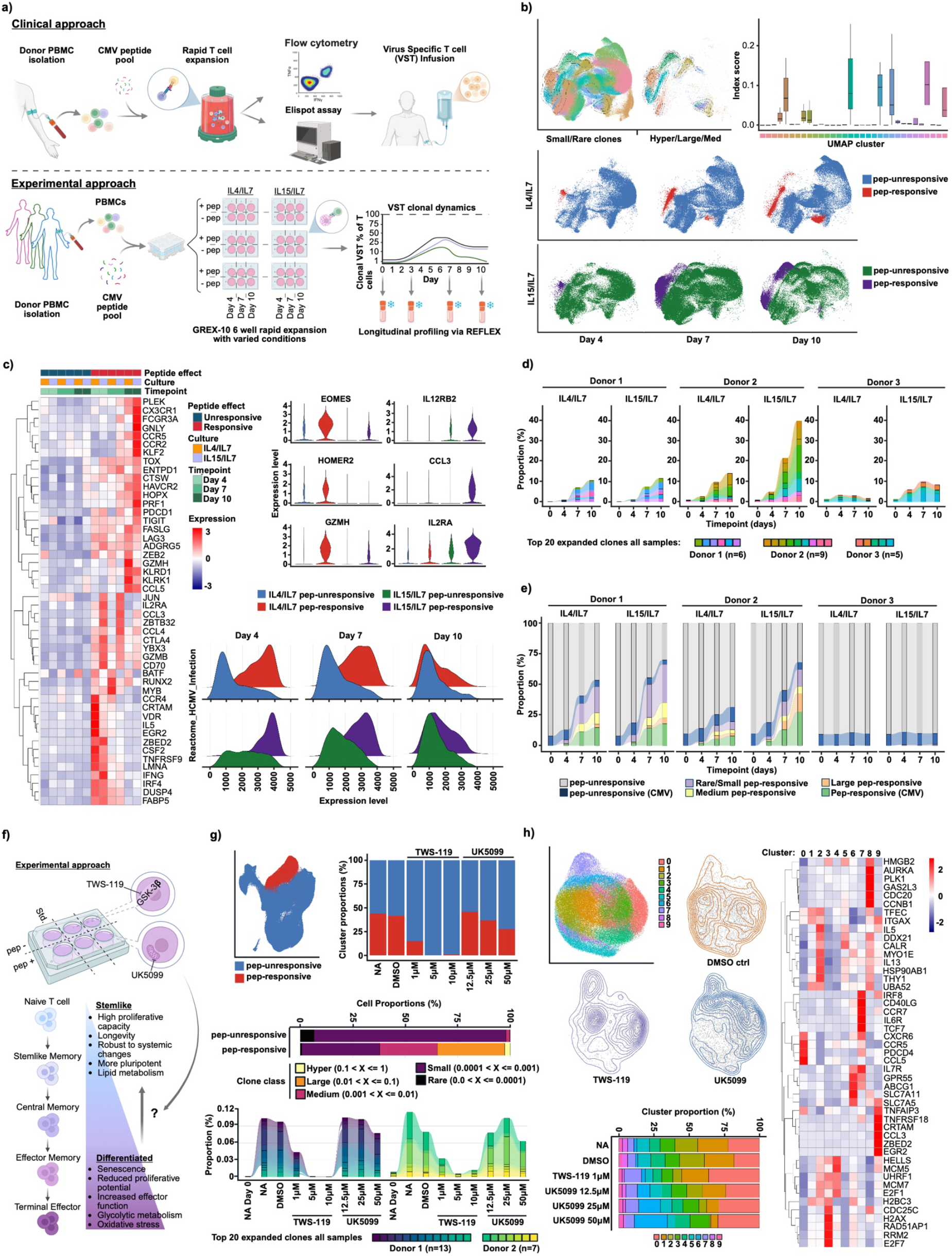
REFLEX allows for comprehensive profiling of VST heterogeneity across donors, cytokines, and metabolic perturbations. a) Schematic comparing VST clinical and experimental approach. b) UMAPs and boxplot of peptide responsive/clonally expanded clusters by size/condition (see Methods). c) Top differentially expressed (DE) (heatmap) and select individual genes (violins) by condition. Ridge plots show pathway enrichment for Reactome_HCMV_infection. d) Expansion of top 20 CDR3 clonotypes and e) expansion of annotated CMV and orphan clones by condition. f) Schematic of VST expansion +/-UK5099 or TWS-119. g) UMAP and bar plot (upper) showing peptide responsive clustering and proportions by treatment. Bar (lower) and alluvial plots depicting clonal expansion by class/size and clonal proportions for top 20 clonotypes by sample. h) UMAP clustering, density plots, and cluster proportions by treatment (bar plot). Heatmap representing top 5-6 DE genes/cluster.

*(Results continued)* UMAP projection identified clear cytokine/peptide dependent evolution of subpopulations over time (Fig2b) which correlated with activation-associated gene expression and HCMV-reactive GSEA (Fig2c). Peptide-expanded TCR subclones were detected in only 2/3 donors, exemplifying the variability seen in primary literature^17^ (Fig2d, SuppFig8a). Of unique peptide expanded CDR3, 4,030 were associated with CMV epitope species (VDJdb, Fig2e), however, >13,000 were orphan, indicating novel identification of CMV responsive clonotypes. Strong subclonal association with CMV specific dCode dextramers (Immudex) further supported this, demonstrating REFLEX compatibility with ADT modalities while highlighting further avenues for discovery using this technology (SuppFig9, Methods).

Developing methods for expansion of VSTs with specific phenotypes/functionalities is of interest to the field. Evidence suggests promoting stem/naïve-like memory T cells could improve clinical efficacy^18^. Targeted metabolic inhibitors including UK5099 (pyruvate transport) and TWS-119 (GSK3β/Wnt) can promote “stemness” during CAR-T generation^19,20^, however, this has not been rigorously explored for VSTs. To demonstrate the multiomic capabilities of REFLEX in pursuing such questions, we expanded VSTs from two CMV reactive donors (above) in the presence of titrated UK5099 or TWS-119 (Fig2f). Both inhibitors, particularly TWS-119, showed reduced peptide reactive expansion (Fig2g), presumably tied to reduced mitotic signature (SuppFig10). Top TCR clonotypes varied between donors but showed general similarity across treatments (Fig2g, SuppFig8b). Subsetting and UMAP plotting of peptide responsive cells demonstrated enrichment of clusters 0 and 7 by treatment with either inhibitor (Fig2h). While cluster 0 was defined by markers associated with CD8 tissue residency/localization, expansion, and effector function (*CXCR6/CCR5/PDCD4/CCL5*)^21,22,23,24^, cluster 7 was enriched for stemness markers *TCF7, IL7R*, and *CCR7*, illustrating phenotypic heterogeneity within treatments (Fig2h)^(25)^. Cluster 6 was uniquely enriched in a dose-dependent manner by UK5099 and contained elevated expression of *IL7R* as well as the cholesterol efflux channel protein, *ABCG1*, with known roles in controlling proliferation and suppressing senescence^26^. Though not within the scope of our study, elevated expression of stemness/memory markers within treatment enriched clusters suggests further profiling of metabolic inhibitors with multiomic technologies like REFLEX has the potential to identify areas of improvement to existing therapeutic approaches.

REFLEX combines cost-efficient, high-throughput, paired-chain single-cell sequencing with multiomic capability, linking rich, high-quality TCR information with transcriptomic context. In doing so it opens the door to comprehensive profiling of T cell immunity in research and clinical settings such as pathogenic responses, autoimmunity, and cancer where challenges of scale and cost have been prohibitive. This will massively accelerate the expansion in quality and scope of available datasets and improve predictive modeling capabilities. Additionally, the underlying chemistry of the targeted in-situ reverse transcription and capture is easily accessible and could be adapted to various other biological applications demanding scalable single cell and/or fixed tissue solutions, including the targeted detection of gene translocations or CRISPR editing. Together these attributes position REFLEX as a transformative tool for unlocking previously inaccessible insights into T cell immune heterogeneity and beyond at unprecedented scale and resolution.

## Methods

### RT and Variable Primer Design

Detailed RT primer design and sequence information, including complementary TCR alpha and beta constant sequences, 10x sample barcode, spacer region and splinting oligo complementarity region can be found in supplemental tables 1-4. Sequences targeting TCR alpha and beta chain constant regions were adapted from Boria 2008, Fahad 2022, Tu 2022^27,28,29^. Variable region primers were adapted from Genolet 2023 and can be found in supplementary table 4. Oligos were synthesized by Integrated DNA Technologies (IDT). Variable primers were purified using standard desalting while splinting oligos and 5’ phosphorylated RT primers were HPLC purified. Concentrated RT probes and sample barcode matched splinting oligos (100 uM each) were then mixed at 30% RT Probe + 45% Bridging Oligo + 25% 10X TBS. Each alpha and beta barcoded TCR RT Probe was then annealed at 95°C for 2 minutes with a ramp down to 4°C at 0.2C/second. Annealed TCR RT probes (alpha and beta, 30 uM each) were then mixed equally by volume for a final concentration of 15 uM per probe barcode. To avoid freeze-thaw cycles, prepared TCRalpa/beta mixtures were aliquoted into single-use volumes and stored at -20°C for later use.

### Custom FLEX Probe Design

Custom FLEX probes for CD45 isoforms were designed adhering to 10x Custom Probe Design Technical Note (CG000621, Rev D)^30^. Probe lists can be found in Supplemental Figure 4.

### VST Culture and Fixation

Culture was performed using GREX-10, 6 well plates (Wilson Wolf), largely following a previously published rapid expansion protocol^15^. Briefly, PBMC were thawed, pelleted at 400 x g and resuspended in 2% BSA/PBS for counting. Cells were pelleted once again and resuspended at 1.5×10^7^ cells per mL in CTL culture media including 300ng/mL each of CMV derived pp65 and IE-1 peptide pools (Peptivator, Milltenyi) or vehicle control (water). CTL culture media contained 44% RPMI (Thermo), 44% Clicks (Sigma), 10% FBS (Thermo), 1% PenStrep (Thermo), 1x Glutamax (Thermo). Cells were incubated/pulsed for 1h at 37C. Following peptide pulse, cells were distributed into GREX-10 6 well plates containing 30mL total volume CTL media. Media was supplemented with IL4 (400U/mL, Thermo), IL7, and/or IL15 (10ng/mL, Thermo) as indicated. On day 7 of culture, the upper 15mL of media was carefully removed from each well without disturbing cells and replaced with 15mL fresh CTL media including 2x cytokine addition. If applicable, treatment with metabolic inhibitors UK5099 (Sigma), TWS-119 (Selleck Chem.) or vehicle/DMSO, was initiated on day 1 of culture. Further addition of inhibitors was done at 1x concentration and occurred only during day 7 media exchange. Individual wells were harvested at indicated timepoints during VST culture. Upon harvest, cells were washed once with cold 0.4% BSA/PBS then once with cold PBS alone prior to pelleting and resuspension in cold 10x fixation buffer per the standard FLEX protocol. Cells were counted and fixed at a maximum of 1×10^7^ per mL fixation buffer^31^. Fixation occurred for 20-26 hours at 4C. After fixation, cells were pelleted at 850 x g and resuspended in 1mL ice cold quenching buffer prior to storage at -80°C with 10x Enhancer and 50% glycerol, per 10x FLEX guidelines (CG000478, Rev D)^31^.

### Cell Lines

Parental Jurkat T cells were originally purchased from ATCC and were transduced with pMW39 lentivirus (CXCR4 sgRNA) + pMW5 lentivirus (dCas9) by our group for use in other, unrelated, studies. No modifications were made to the native Jurkat TCR. The Raji CCL-86 cell line was purchased from ATCC.

### Donor Sample Acquisition

PBMC samples were sourced commercially via Bloodworks and Stemcell technologies. HLA typing was provided by source company (SuppTable4).

### Fluorescent and Dextramer Staining

After cell culture, samples for dextramer and flow staining were aliquoted at a concentration of 5*10e6 cells/mL in 1mL in 12x75mm polypropylene round bottom test tubes (VWR, 60818-576). Samples were spun down at 400 x g (4°C) and resuspended in 100 μL 2% FBS in PBS to be transferred to a round bottom 96 well plate (Falcon Corning, 351177). After transfer, cells were spun down once again to be resuspended in a dextramer staining mix. To create this dextramers were spun down at 10,000 xg (4°C) for 10 minutes and a bulk staining mix containing dextramer (Supplemental Tables 5 and 6) and 2% FBS in PBS was made using the recommended titer provided by Immudex. Cells were stained for 1 hour at 4°C in the dark followed by a 150 μL wash with DPBS without calcium or magnesium (Corning, 21-031-CM). For transition straight into further CITE staining and REFLEX, see the following methods section: REFLEX assay (CITE). For analysis by flow cytometry, samples were centrifuged at 750 x g for 5 min (4°C) and resuspended in a viability and blocking mix made up of Fixable Viability Dye 510 (BD biosciences), Human TruStain FcX (BioLegend), and DPBS (Corning). Cells were incubated at 4°C in the dark for 30 min. After incubation, cells were diluted with 150 μL of cell staining buffer and centrifuged at 75 x g at 4°C for 5 min to pellet. Supernatants were removed and cells resuspended in a cocktail of fluorophore conjugated antibodies (SuppTable6) then incubated at 4°C in the dark for 30 min. Post incubation, samples were again diluted, spun down and washed twice in 200 μL of cell staining buffer. Samples destined for processing and analysis only were resuspended in a final volume of 250 μL of cell staining buffer for acquisition on a Cytek Aurora spectral cytometer.

### REFLEX Assay (Thaw)

Cryovials of PBMCs or cell lines were thawed in a 37°C water bath for 2 minutes before being transferred into 40 mL of pre-warmed AIM V media (37°C) (GIBCO), inverted 3 times, and centrifuged for 5 minutes at 400 x g (4°C) in a swinging bucket rotor. Supernatant was removed and pelleted cells were washed with 25 mL of cold (4°C) PBS + 2% BSA, then centrifuged for 5 minutes at 400 x g (4°C) in a swinging bucket rotor. Pelleted cells were then resuspended in 1 mL PBS (4°C) and 27 μL was removed for counting. The counting fraction was stained 1:1 with 27 μL of AO/PI solution and counted on a Nexcelom Cellaca MX cell counter. Sample viability of 90% or greater was targeted.

### REFLEX Assay (CITE)

Dextramer staining was performed as described above and in accordance with Immudex dCODE Dextramer (10X) Package protocol (Immudex)^32^. Following dextramer staining and washing, cells were stained and washed using BioLegend’s TotalSeq™-C TBNK cocktail (Cat. No 399903, SuppTable5) according to the Cell Surface & Intracellular Protein Labeling for Chromium Fixed RNA Profiling protocol (CG000529, Rev C)^32^.

### REFLEX Assay (Fixation)

PBMCs or cultured T cells with or without prior CITE antibody or dextramer staining, were fixed according to the Fixation of Cells & Nuclei for Chromium Fixed RNA Profiling protocol (CG000478, Rev D)^31^ in a final concentration of 4% Paraformaldehyde for at 20°C for 1 hour or at 4°C overnight (16-24 hours). Fixed samples were subsequently quenched following the 10x protocol above and either advanced immediately into hybridization or stored at -80°C following 10x guidelines.

### REFLEX Assay (Hybridization)

Up to 500,000 fixed cells per sample barcode were hybridized/barcoded for multiplexing using 10x Genomics Fixed RNA Feature Barcode Multiplexing Kit (PN-1000628). Probe hybridization was completed according to the 10x Genomics protocol for Multiplexed samples with Feature Barcode technology for Protein using Barcode Oligo Capture (CG000673, Rev B)^32^. An additional 6.7 μL of custom TCR RT probes were spiked in (1 μM/probe). Where applicable, custom GEX probes for CD45 were spiked in at 800 nM/probe in a total volume addition of 5 μL. Hybridization occurred for 16-24 hours at 42°C.

### REFLEX Assay (Post-Hybridization Washes)

Following hybridization, excess probes were removed with 3 cycles of washing in 10x Post-Hyb Wash buffer [5% Concentrated Post-Hyb Wash Buffer and 5% Enhancer by volume in molecular biology grade water (MBGW)]. This was done either manually or using an automated Tecan liquid handler. A total volume of 115 μL of hybridized cells per barcoded sample were transferred to separate 1.5 mL Eppendorf tubes. Each well containing barcoded cells (strip tube or 96-well plate) was rinsed with 175 μL wash buffer to maximize cell retention. Individual sample tubes were volume normalized to 1 mL by adding 710 μL of Post-Hyb Wash buffer. Cells were incubated for 10 minutes at 42°C, then pelleted for 5 min at 850 x g. Supernatants were then aspirated, and pellets were resuspended in 400 μL wash buffer and then incubated for 10 min once again. These steps were repeated a total of 3 x 10 minute incubations. After final pelleting, 600 μL of remaining supernatant was removed, and the pellet resuspended in 80 μL post-hyb resuspension buffer [5% concentrated post-hyb wash buffer by volume]. Samples were transferred to a 96 well plate and spun at 850g to pellet again. Supernatant was removed for a final volume 50.8 μL which was used to gently pipette resuspend cell pellets.

### REFLEX Assay (Ligation)

GEM master mix, containing 60% 10x GEM reagent mix (10x, PN 20000491) and 5% 10X Reducing Agent B (10x, PN 2000087) by volume in MBGW was prepared for use in ligation and reverse transcription reactions. Ligation master mix consisted of 33.3 μL GEM master mix, 5 μL ATP (New England Biolabs, P0756L), 7.5 μL SplintR ligase [New England Biolabs, M0375L, 25,000 u/mL], and 3.3 μL RNase Inhibitor [Ambion, Superase-In, AM2696 20U/μL]. 49.2 μL ligation master mix was added to each cell suspension for a final volume of 100 μL. Ligation reactions were incubated for 60 min at 25°C after which cells were pelleted at 850 x g for 5 minutes and supernatants removed. Pellets were washed in 95 μL chilled post-hyb resuspension buffer, pelleted, and 60 μL supernatant removed, leaving 40 μL final volume. Cell pellets were gently resuspended in the remaining volume.

### REFLEX Assay (Reverse Transcription)

RT master mix consisted of 33.34 μL GEM master mix as described above, along with 5 μL Recombinant Albumin (NEB, B9200S, 20mg/mL), 5 μL dNTPs (NEB, N0447L 10mM), 3.3 μL 0.1 M dithiothreitol, 3.3 μL Superase-IN (see above), and 10 μL ProtoScript II RT enzyme (New England Biolabs, M0368L). 60 μL of RT master mix per sample/well was added to pre-existing 40 μL of volume for a final volume of 100 μL. Samples were then incubated for 45 min at 45°C. After incubation, 120 μL post-hyb resuspension buffer were added to samples which were then mixed by gentle pipetting and kept on ice.

### REFLEX Assay (GEM Generation)

Cells were counted on a Nexcelom Cellaca MX. Samples were then pooled at equivalent cell concentrations, washed in Post-Hyb resuspension buffer, and strained using 30 μM strainers (Miltenyi, 130-041-407) as noted in the 10x User Guide CG000673, Rev B. The single cell suspension pool was loaded following 10x guidelines onto Chip Q for GEM generation at a concentration of 16,000 cells per barcoded sample or 256,000 cells for 16-plex pool. GEMs were transferred to strip tubes and incubated following standard 10x guidelines to allow for in-GEM ligation and cell barcoding.

### REFLEX Assay (Library Prep, Pre-Amp)

Following GEM incubation, 125 μL of Recovery Agent were added to break the GEM emulsion. Mixtures were inverted 10 times and incubated for 2 minutes at room temperature. 125 μL of recovery agent was then carefully removed from the bottom of each well. The remaining approximately 65 μL of recovered GEM sample was combined with a master mix containing 27 μL of Amp Mix (10x PN-2000047), 10 μL of Pre-Amp Primers B (10x PN-20000529), 1.5 μL of TruSeq1 [1 μM], and 3 μL of a pre-mixed TCR Variable Primer Pool (See Supplemental Table 3) [0.66 μM] for a total volume of 106.5 μL of master mix plus recovered GEM. Samples were inverted 10 times to mix and placed in a thermal cycler with the following cycling conditions: 98C for 3 minutes; 8 cycles of 98C/20 seconds, 60C for 30 seconds, 72C for 2 minutes; 72C for 5 minutes, 4C infinite hold. Upon completion of the PCR, tubes were pulsed briefly on a minifuge to collect liquid. 90 μL were then transferred into a new 1.5 mL Eppendorf tube. A 1.8X SPRI bead cleanup was performed and sample was eluted into 100 μL of elution solution in accordance with 10x User Guide CG000673 Rev B.

### REFLEX Assay (Library Prep, Re-Amplification)

Pre-amplified library eluates were split into two or three pools for preparation of gene expression, TCR, and ADT libraries (when applicable). Gene expression and ADT libraries were prepared using 20uL pre-amp product and following 10x guidelines for Multiplexed samples with Feature Barcode technology for Protein using Barcode Oligo Capture (CG000673, Rev B)^32^. Linear PCR master mix for TCR libraries consisted of 20 μL 1X KAPA HiFi HotStart (Roche, KK260), 10 uL molecular biology grade water (Fisher, BP2819100), and 0.5 μL TruSeq2 primer at 123.4 nM per reaction. TCR libraries utilized 40 μL of pre-amp product total, divided into 4, 10 μL aliquots. Each 10 μL aliquot was combined with 30.5 μL master mix in a PCR strip tube and incubated with the following PCR program: 95C/3min, [98C/20s, 62C/30s, 72C/90s] repeat x4 for 5 total cycles, followed by a 72C/5min final extension, hold 4C. After the 5-cycle linear PCR was completed, 0.5 μL TruSeq1 primer, at 123.4 nM per reaction, was spiked into each reaction and an exponential PCR was performed with the following conditions: 95C/3min, [98C/20s, 62C/30s, 72C/90s] repeated x7 for 8 total cycles, followed by a 72C/5min final extension, hold 4C. Following exponential PCR, each of the four 41 μL reactions per library condition were repooled for a total volume of 164 μL. Samples were SPRI cleaned at 0.8X ratio, eluted in 50 μL aforementioned “elution solution”.

### REFLEX Assay (Indexing TCR Libraries)

PCR indexing master mix was prepared with 50 μL 1X Kapa HiFi HotStart and 10 μL molecular biology grade water per reaction. Indexing reactions combined 20 μL of cleaned re-amplified product, 60 μL master mix, and 20 μL Dual Index TT Set A (10X Genomics, PN-1000215). Each library used a unique TT index combination enabling later identification via demultiplexing. Samples were inverted to mix and briefly spun down. Thermocycler conditions used were as follows: 98C/45s, [98C/20s, 54C/30s, 72C/90s] repeated x12-15 for 13-16 total cycles, followed by a 72C/60s final extension, hold 4C. Cycle number was adjusted to account for cell input during GEM generation. A final SPRIselect bead clean up was performed at a 0.7X ratio with cleaned product eluted into 40 μL Buffer EB (Qiagen, Cat No. 19086).

### REFLEX Assay (Indexing GEX libraries)

GEX library preparation was performed separately per 10x User Guide CG000673 Rev B, Sample Index PCR guidelines, Section 4.1, using 20 μL pre-amplification product with 60 μL Sample Index PCR Mix (Amp Mix PN-2000103, Water) and 20 μL of a unique well of Dual Index TS Set A primers (PN-3000511). The combined solution was gently mixed and spun down. PCR performed per the following guidelines: 98C/45s, [98C/20s, 54C/30s anneal, 72C/20s elongate] repeat 7-14x for a total 8 to 15 cycles, 72C/60s final extension, hold 4C. Cycle number was adjusted to account for cell input during GEM generation. Indexed products were cleaned using a 1.0X SPRIselect bead cleanup and eluted in 40 μL Buffer EB.

### REFLEX Assay (Indexing ADT libraries)

ADT library preparation was performed separately per 10x User Guide CG000673 Rev B, Sample Index PCR guidelines, Section 5.1, using 20 μL pre-amplification product with 60 μL Sample Index PCR Mix (Amp Mix PN-2000103, Water) and 20 μL of a unique index combination from Dual Index TN Set A primers for the ADT libraries (10x, PN 1000250). The combined solution was gently mixed and spun down. PCR performed per the following guidelines: 98C/45s, [98C/20s, 54C/30s, 72C/20s] repeat 7-14x for a total 8 to 15 cycles followed by a 72C/60s final extension, hold 4C. Cycle number was adjusted to account for cell input during GEM generation. Indexed products were cleaned using a 1.0X SPRIselect bead cleanup and eluted in 40 μL Buffer EB.

### Sequencing

scRNAseq libraries were sequenced using a NovaSeq X 25B PE300 or NextSeq2000 P4 300 cycle flow cell, Read1:80 cycles Read2:200 cycles. A read depth of 5000 reads per cell for TCR libraries or 10,000 reads per cell for GEX libraries was targeted. ADT and CITEseq libraries were targeted at 5000 reads per cell.

### Data Analysis

Following sequencing, GEX FASTQ files were processed via Cell Ranger (10X Genomics, v9.0.1). Doublet detection was performed on the resulting cell by gene matrices using Scrublet (v5.0.1) ^33^. Cells were annotated using CellTypist ^3,3^ at 3 resolutions of granularity with the reference as described in Gong et al ^36^. The labeled h5 files were then combined into a single Seurat^37,38,39,40^ object and filtered to remove doublets and dying cells based on Scrublet calls, genes per cell and UMIs per cell.

TCR alignment was performed using MIXCR’s (v4.7) ^41^ “generic-ht-single-cell-amplicon workflow”. Sample barcodes were provided using the –-sample-sheet parameter to demultiplex the TCR libraries. Cell barcode alignment used the 10X barcode whitelist in the –set-whitelist parameter. The resulting clones.tsv files were integrated into the Seurat object using scRepertoire (v2.5.3).

Gene expression analysis utilized standard Seurat object processing functions (NormalizeData, FindVariableFeatures, ScaleData, RunPCA, RunUMAP, FindNeighbors, FindClusters). Differential gene expression analysis used Seurat’s FindMarkers function using a Wilcoxon rank sum test. Escape (v2.5.5) was used to run ssGSEA on the Seurat object using REACTOME’s HCMV Infection pathway. ADT and WNN analysis used standard built-in Seurat workflows. To identify dextramer positive cells, we established a normalized expression cutoff of 1.5, where there was clear separation between background and dextramer high cells (SuppFig9b).

scRepertoire (v2.5.3) was used to analyze the TCR data including calculating clone size, STARTRAC-based clonal expansion index, and generating CDR3 alluvial plots. Captured CDR3 sequences were aligned with VDJdb (9/25/25 release) ^42,43^ to determine antigen epitopes and known pathogens. Levenstein distance was used to calculate the number of mismatches to the closest related CDR3s in the database compared to captured CDR3s.

## Data availability

Raw fastq files from FLEX gene expression and TCR sequencing have been deemed exempt from data protection/privacy laws as they do not contain donor identifiable information. They will be available upon completion of submission to Gene Expression Omnibus (GEO) (delayed due to government shutdown).

## Code availability

Analytical code is available at the following GitHub link: https://github.com/aifimmunology/reflex_manuscript

## Lead contact

Requests for further information or inquiry about resources and reagents should be directed to the lead contact, Peter Skene.

## Acknowledgements

We would like to thank the Allen Institute founder, P.G. Allen, for his vision, encouragement and support. We are additionally thankful to members of the Allen Institute for Immunology, particularly members of the facilities and operations, flow cytometry, bioinformatics operations and automation teams for their support and contributions to maintaining a productive research environment. We thank Claire Gustafson and Maia Boehm for their constructive feedback during manuscript preparation.

## Author contributions

*Conceptualization*: P.J.S., M.R.H., Z.J.T., N.S.K., S.I., T.R.T., *Methodology and experiments*: all authors, *Investigation and data analysis*: Z.J.T., M.R.H., N.S.K., T.J.S., *Visualization*: Z.J.T, M.R.H., N.S.K., S.I., T.J.S., *Writing - original draft*: M.R.H., *Review and editing*: all authors

