## Supplemental figures and tables for "REFLEX, a Novel Immune Profiling Assay, Combining TCR Repertoire and Multiome at Massively Scalable Single-cell Resolution to Catapult Exploration of T-cell Derived Immunity"

*Supplemental Figures/Data*

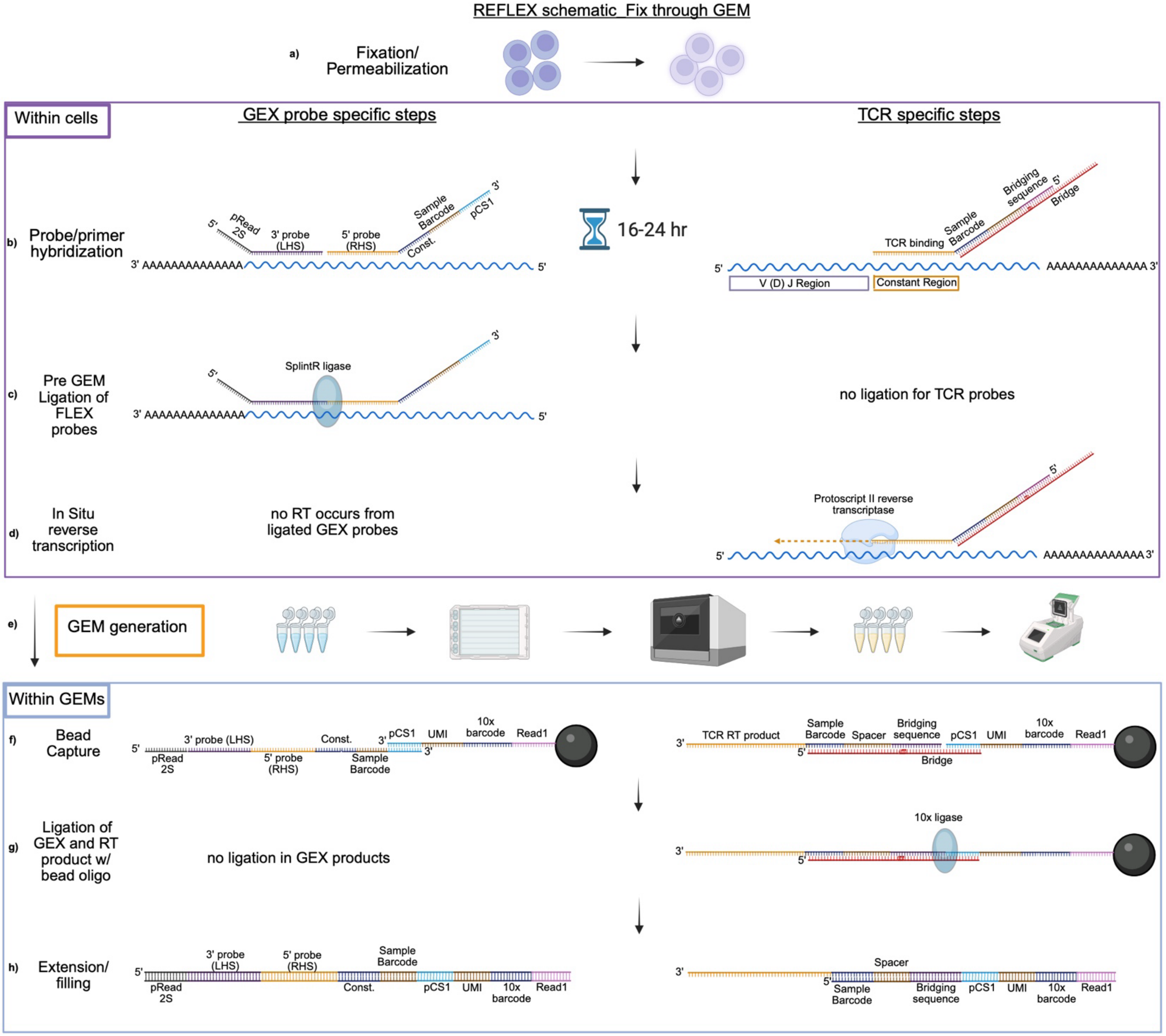

**Supplemental Figure 1. Detailed step-by-step schematic of REFLEX chemistry leading up to emulsion**. Showing representative primer/probe oligos and relative orientations. GEX and TCR pictured separately but with various steps occurring in parallel. a) Cell fixation and permeabilization. b) Overnight hybridization of 10X FLEX probes and pre-annealed bridge/TCR constant region binding RT primers. c) Ligation of left and right hand 10X probes to prevent off-target/strand displacing RT. d) Reverse transcription of TCR CDR3 variable region. e) Pooling, filtering, counting, and 10X GEM generation on Chromium platform. f) Bead capture of FLEX probes and TCR RT products by GEM beads in emulsion. g) Ligation of GEX and RT products to GEM beads, ligation of RT product eliminating bridging sequence dependency, h) extension and gap-filling of GEX and RT products.

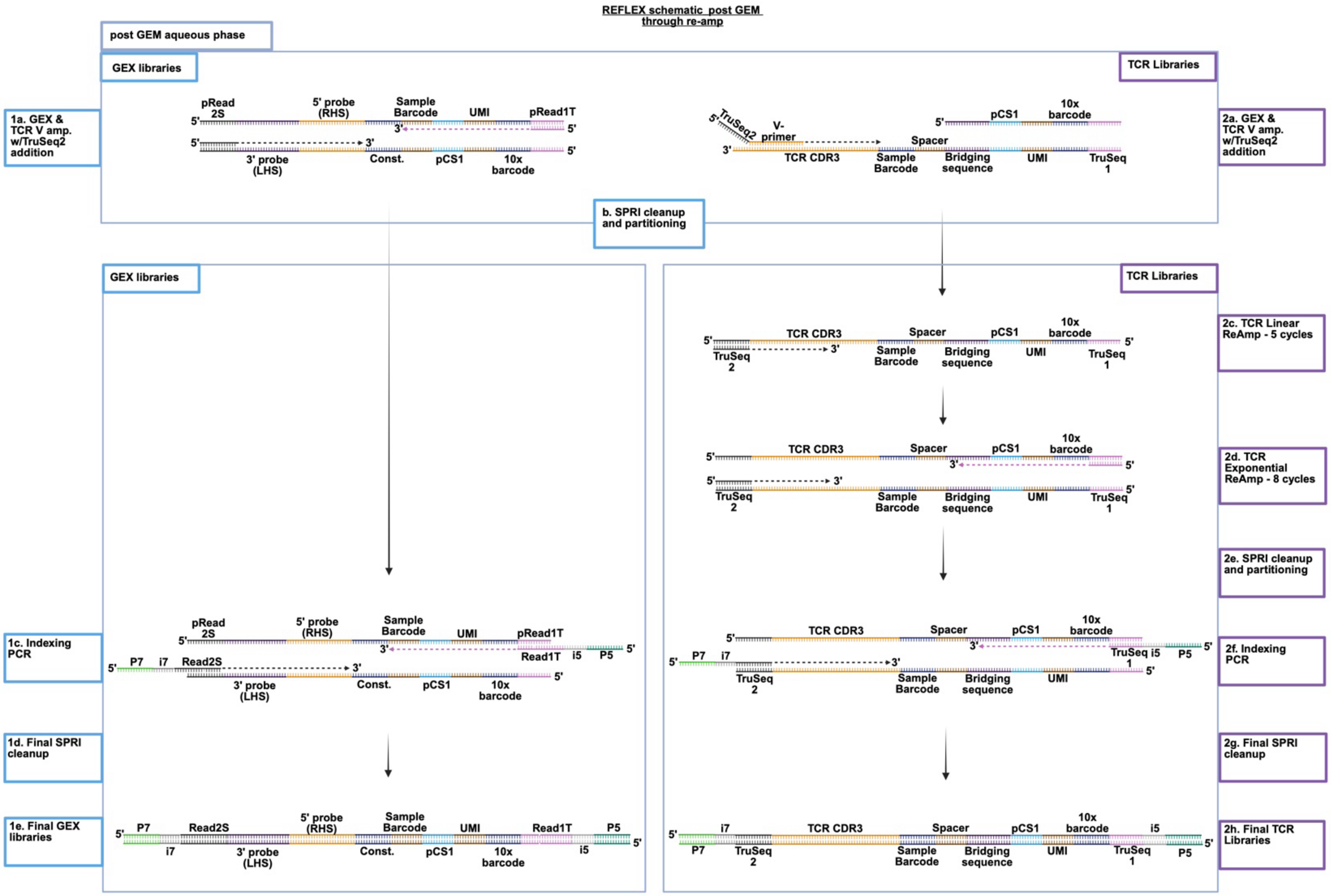

**Supplemental Figure 2. Detailed schematic of REFLEX chemistry post GEM.** Showing representative primer/probe oligos and relative orientations. GEX portion shown with 1a) combined preamplification after GEM emulsion disruption, including pre-amp primer group B (or C if library contains ADTs) and addition of TruSeq 1 primers, b) 1.8X SPRI clean up and partitioning of pre-amp product, 1c) indexing PCR, addition of Illumina-compatible p5/p7 primers from 10X Dual Index TS kit (or TN for ADT's), 1d) 1X SPRI clean up, 1e) fully indexed GEX (or ADT) product structure, including sequencing primers, 10X barcode, and UMI for sample identification. TCR portion shown with 2a) combined reamplification after GEM emulsion disruption, TCR-variable pool and TruSeq1 spike-in primers for TCR capture and amplification, 2b) 1.8X SPRI clean-up and partitioning of pre-amp product, 2c) linear amplification phase of TCR products, using only TruSeq2 primer to preferentially amplify TCR libraries for 5 cycles, 2d) TruSeq1 primer added to re-amplification for 8-cycle exponential amplification of TCR libraries, 2e) pooling of reamplification products and 0.8X SPRI clean up, 2f) indexing PCR, addition of Illumina-sequencing compatible p5/p7 primers from 10X Dual Index TT kit, 2g) 0.7X SPRI clean up, 2h) fully indexed TCR product structure, including sequencing primers, 10X barcode, and UMI for sample identification.

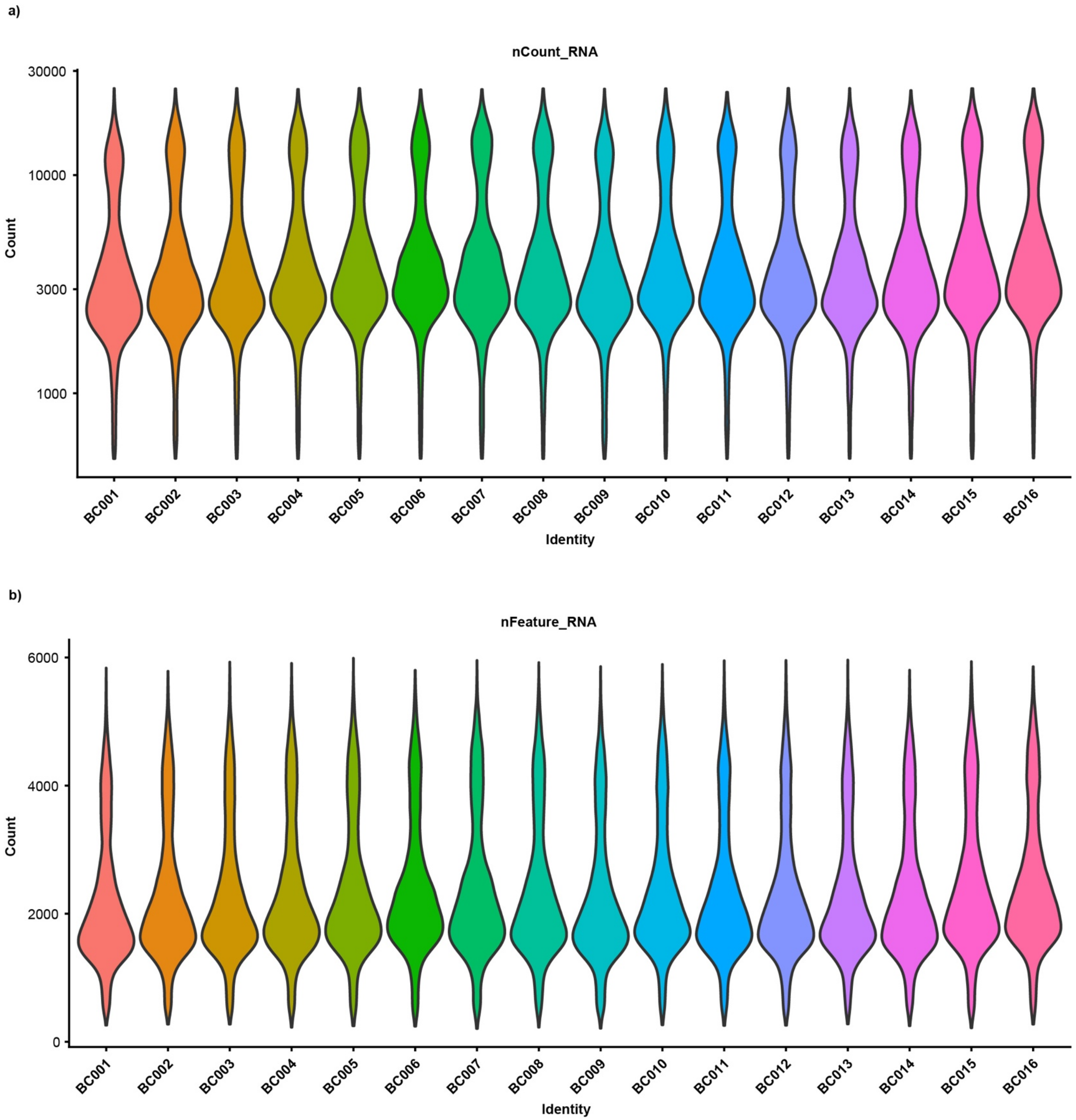

**Supplemental Figure 3. REFLEX gives consistent performance across multiplex barcodes.** a) UMI and b) features detected across 16 multiplexed barcodes in PBMCs.

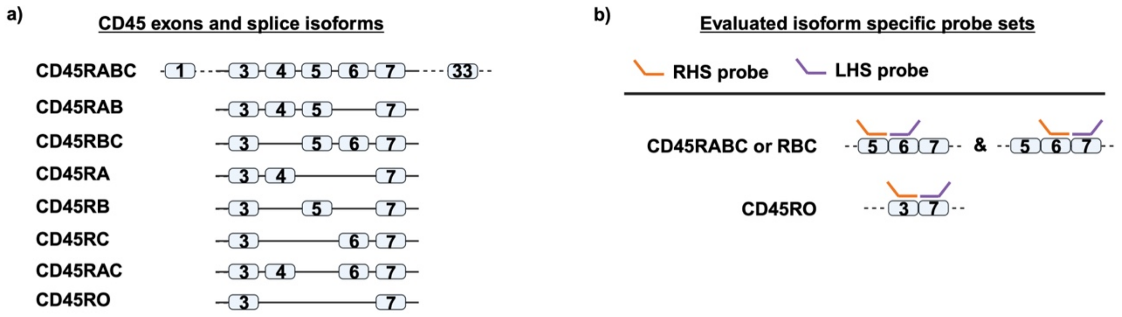

c)

| gene_id | probe_seq | probe_id | included | region | gene_name |
|---|---|---|---|---|---|
| ENSG00000081237 | GTTTCAGAGGCATTAAGGTAGGCATCAGTGGGGAAGGTGTTGGGCTTTG | ENSG00000081237\|PTPRC\|E3_7jxn | TRUE | spliced | PTPRC |
| ENSG00000081237 | GCTGTACTCCTCTCTCCTGGGACATCTGAGATAGCATTGCTGCCTGGGGT | ENSG00000081237\|PTPRC\|E5_6jxn | TRUE | spliced | PTPRC |
| ENSG00000081237 | GTTTCAGAGGCATTAAGGTAGGCATCTGAGGTGTTCGCTGTGATGGTGGT | ENSG00000081237\|PTPRC\|E6_7jxn | TRUE | spliced | PTPRC |

**Supplemental Figure 4. REFLEX is compatible with custom FLEX probes for CD45 isoforms.** a) CD45 isoform exon inclusion/exclusion, b) Schematic design for tested isoform specific probe pairs and c) target sequence information for combined left and right-hand side probes (not inclusive of overhangs).

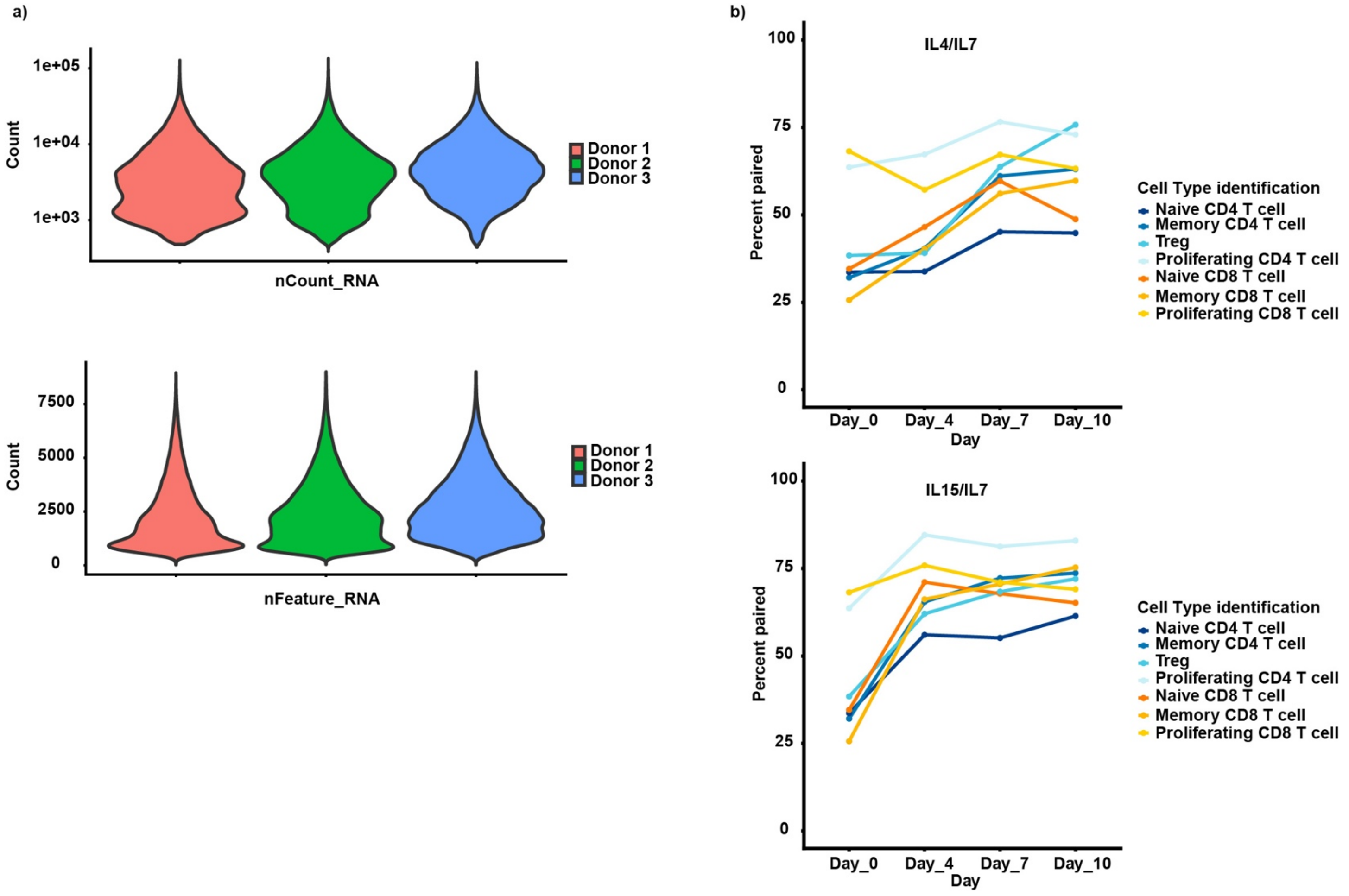

**Supplemental Figure 5. REFLEX allows for gene expression profiling from fixed/stored samples.** a) GEX performance by donor and b) TCR performance over time and by T cell subset ID (donors combined).

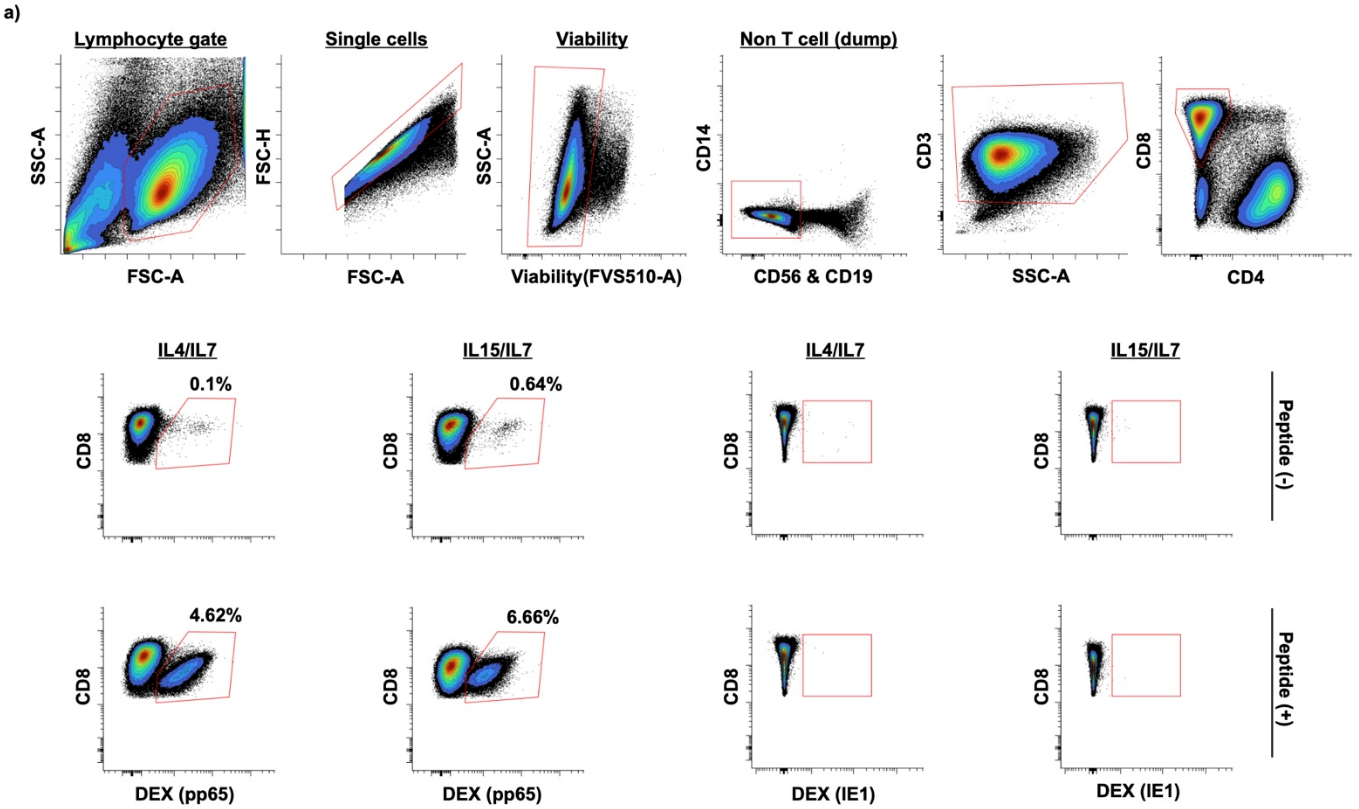

**Supplemental Figure 6. CMV peptide pulse and rapid expansion successfully generated VSTs.** a) Gating scheme and flow cytometry confirmation of CMV reactive T cell expansion by fluorophore conjugated dextramer stain in a single representative donor.

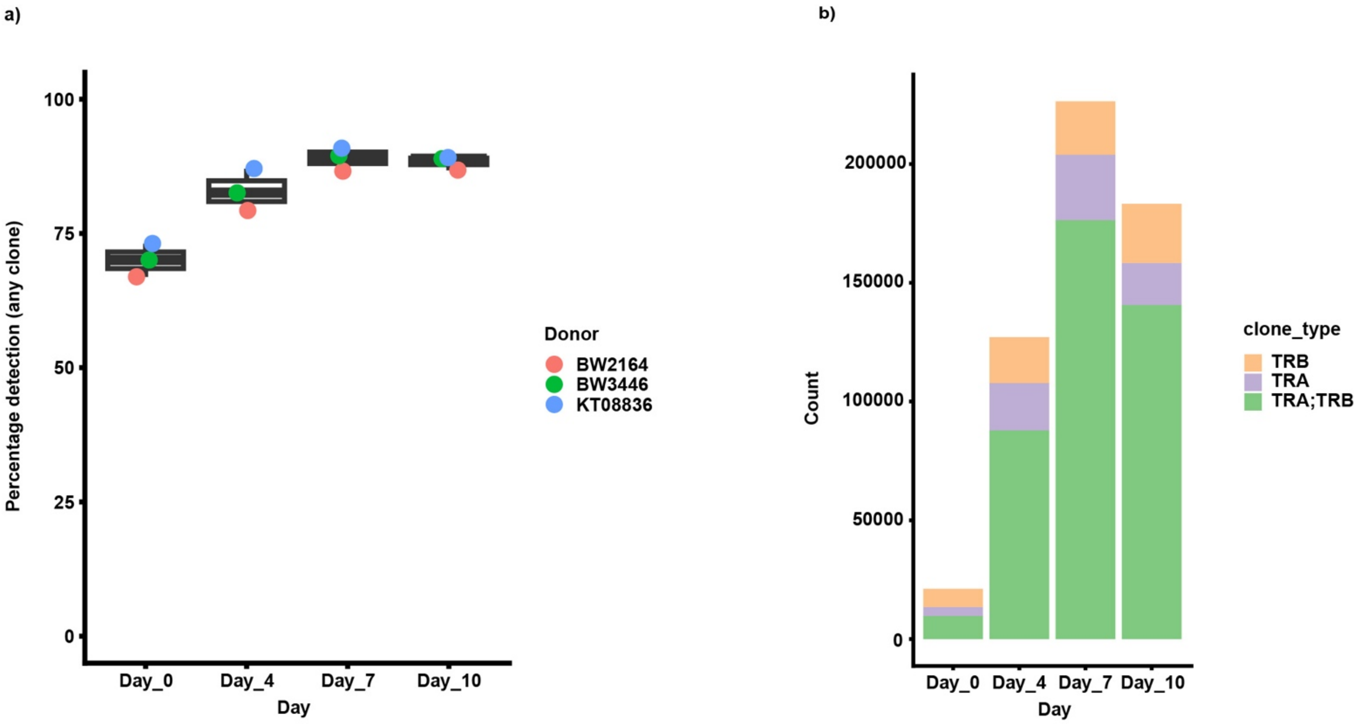

**Supplemental Figure 7. REFLEX shows strong detection of TCR clones across donors and over time.** a) TCR “any” clone detection frequency per cell by time point and b) total number of paired or alpha/beta single clones detected by timepoint.

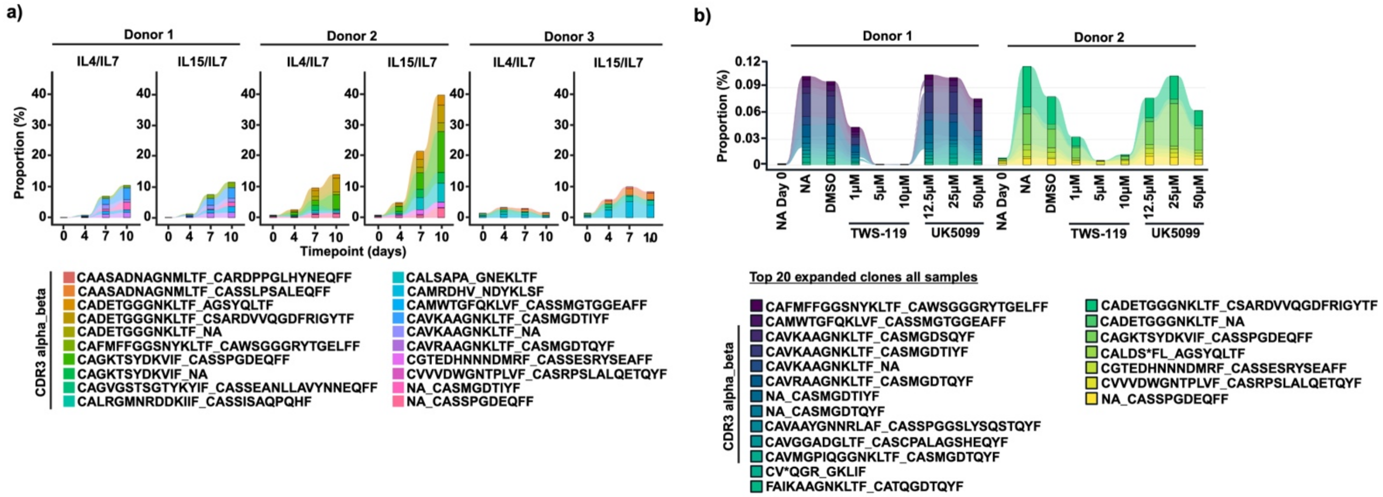

**Supplemental Figure 8. REFLEX tracks clonal expansion across culture timepoints with and without metabolic perturbation.** a) CDR3 amino acid sequences from Figure 2d and b) Figure 2g.

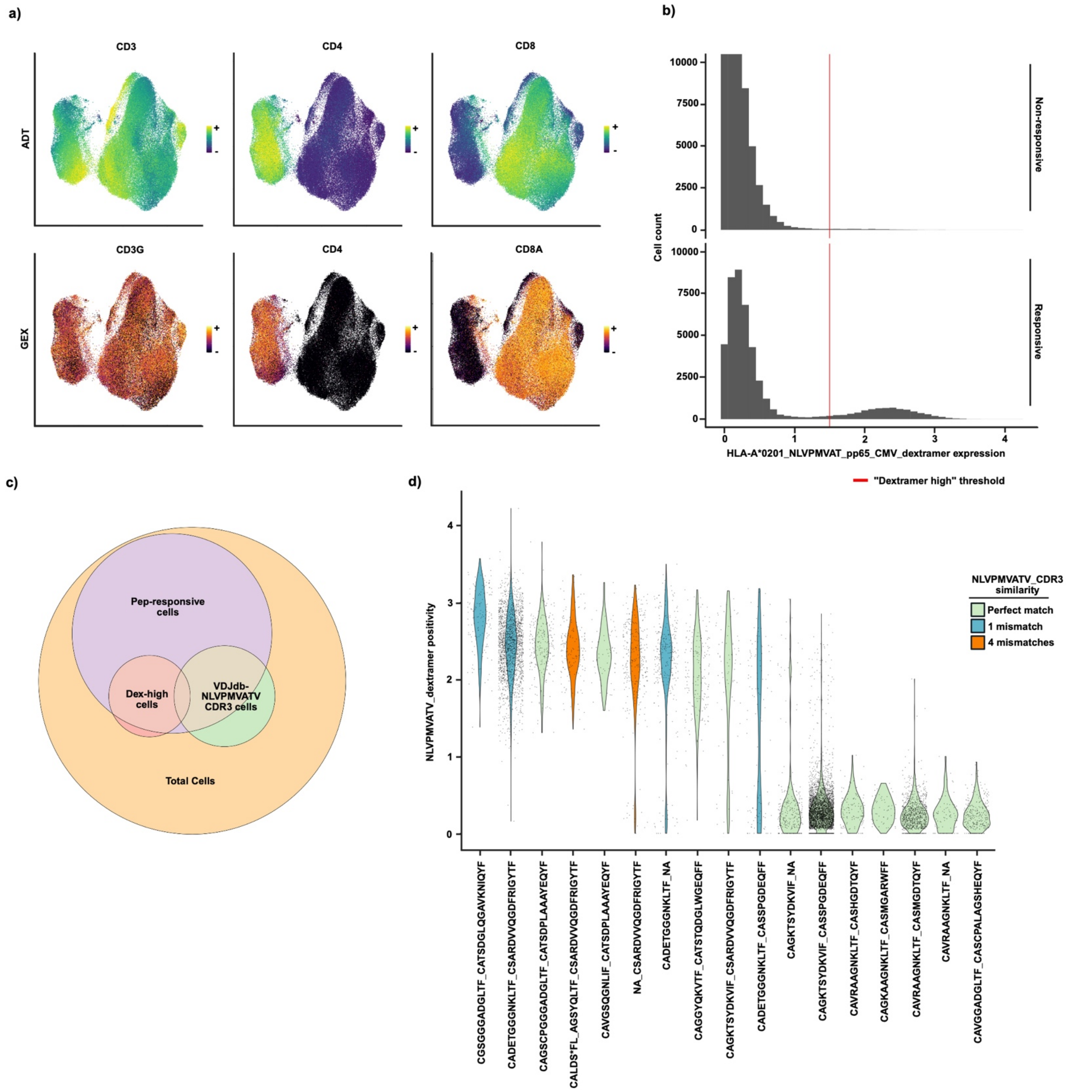

**Supplemental Figure 9. ADT and Dextramer staining identify cell types and potentially novel NLVPMVATV responsive TCRs through REFLEX.** T cell labeling with Total Seq-C TBNK and oligo conjugated “dCode” HLA-A*0201 + NLVPMVATV peptide dextramers demonstrates REFLEX compatibility with ADT modalities and supports novel identification of NLVPMVATV peptide reactive CDR3 a/b pairs. a) UMAPs showing select T cell markers from Total Seq TBNK panel (Biolegend) and corresponding GEX. b) Selection of threshold for “Dextramer high” cells. c) Venn diagram showing overlap between total, peptide responsive, dextramer high, and CDR3 pairings associated with NLVPMVATV peptide by VDJdb. d) Violin plots showing overlap between top 10 dextramer high and top 10 expanded CDR3 from the dataset. Colors indicate similarity between sequenced CDR3 and database.

a)

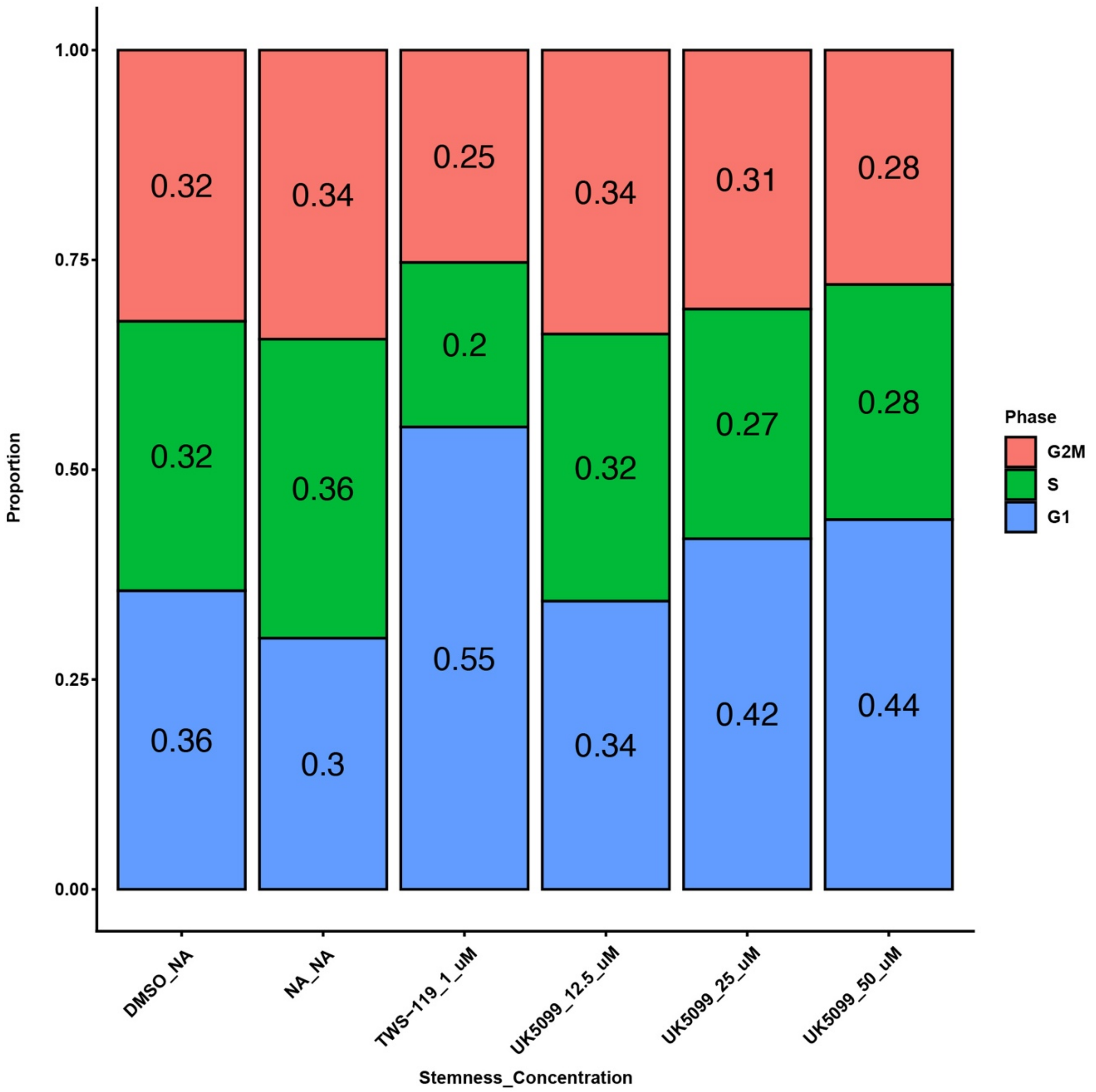

**Supplemental Figure 10. REFLEX profiles effects of metabolic perturbations on cell cycle phase.** Cell cycle phase proportions based on gene expression signature across treatments (showing all cells/donors).

| Name | Target Region Primer Sequence | Purification |
|---|---|---|
| TRAC-RT-BC001-1-P | /5phos/ctagcgtacgagcactccagattacttcca**ACTTTAGG**CGGTGAATAGGCAGACAGACTTGTCACTGG | HPLC |
| TRBC-RT-BC001-1-P | /5phos/ctagcgtacgagcactccagattacttcca**ACTTTAGG**ACCAGTGTGGCCTTTTGGGTGTGGGAG | HPLC |
| TRAC-RT-BC002-1-P | /5phos/ctagcgtacgagcactccagattacttcca**AACGGGAA**CGGTGAATAGGCAGACAGACTTGTCACTGG | HPLC |
| TRBC-RT-BC002-1-P | /5phos/ctagcgtacgagcactccagattacttcca**AACGGGAA**ACCAGTGTGGCCTTTTGGGTGTGGGAG | HPLC |
| TRAC-RT-BC003-1-P | /5phos/ctagcgtacgagcactccagattacttcca**AGTAGGCT**CGGTGAATAGGCAGACAGACTTGTCACTGG | HPLC |
| TRBC-RT-BC003-1-P | /5phos/ctagcgtacgagcactccagattacttcca**AGTAGGCT**ACCAGTGTGGCCTTTTGGGTGTGGGAG | HPLC |
| TRAC-RT-BC004-1-P | /5phos/ctagcgtacgagcactccagattacttcca**ATGTTGAC**CGGTGAATAGGCAGACAGACTTGTCACTGG | HPLC |
| TRBC-RT-BC004-1-P | /5phos/ctagcgtacgagcactccagattacttcca**ATGTTGAC**ACCAGTGTGGCCTTTTGGGTGTGGGAG | HPLC |
| TRAC-RT-BC005-1-P | /5phos/ctagcgtacgagcactccagattacttcca**ACAGACCT**CGGTGAATAGGCAGACAGACTTGTCACTGG | HPLC |
| TRBC-RT-BC005-1-P | /5phos/ctagcgtacgagcactccagattacttcca**ACAGACCT**ACCAGTGTGGCCTTTTGGGTGTGGGAG | HPLC |
| TRAC-RT-BC006-1-P | /5phos/ctagcgtacgagcactccagattacttcca**ATCCCAAC**CGGTGAATAGGCAGACAGACTTGTCACTGG | HPLC |
| TRBC-RT-BC006-1-P | /5phos/ctagcgtacgagcactccagattacttcca**ATCCCAAC**ACCAGTGTGGCCTTTTGGGTGTGGGAG | HPLC |
| TRAC-RT-BC007-1-P | /5phos/ctagcgtacgagcactccagattacttcca**AAGTAGAG**CGGTGAATAGGCAGACAGACTTGTCACTGG | HPLC |
| TRBC-RT-BC007-1-P | /5phos/ctagcgtacgagcactccagattacttcca**AAGTAGAG**ACCAGTGTGGCCTTTTGGGTGTGGGAG | HPLC |
| TRAC-RT-BC008-1-P | /5phos/ctagcgtacgagcactccagattacttcca**AGCTGTGA**CGGTGAATAGGCAGACAGACTTGTCACTGG | HPLC |
| TRBC-RT-BC008-1-P | /5phos/ctagcgtacgagcactccagattacttcca**AGCTGTGA**ACCAGTGTGGCCTTTTGGGTGTGGGAG | HPLC |
| TRAC-RT-BC009-1-P | /5phos/ctagcgtacgagcactccagattacttcca**ACAGTCTG**CGGTGAATAGGCAGACAGACTTGTCACTGG | HPLC |
| TRBC-RT-BC009-1-P | /5phos/ctagcgtacgagcactccagattacttcca**ACAGTCTG**ACCAGTGTGGCCTTTTGGGTGTGGGAG | HPLC |
| TRAC-RT-BC010-1-P | /5phos/ctagcgtacgagcactccagattacttcca**AGTGAGTG**CGGTGAATAGGCAGACAGACTTGTCACTGG | HPLC |
| TRBC-RT-BC010-1-P | /5phos/ctagcgtacgagcactccagattacttcca**AGTGAGTG**ACC | HPLC |

| | | |
|---|---|---|
| | AGTGTGGCCTTTTGGGTGTGGGAG | |
| TRAC-RT-BC011-1-P | /5phos/ctagcgtacgagcactccagattacttcca**AGAGGCAA**CGG TGAATAGGCAGACAGACTTGTCACTGG | HPLC |
| TRBC-RT-BC011-1-P | /5phos/ctagcgtacgagcactccagattacttcca**AGAGGCAA**ACC AGTGTGGCCTTTTGGGTGTGGGAG | HPLC |
| TRAC-RT-BC012-1-P | /5phos/ctagcgtacgagcactccagattacttcca**ACTACTCA**CGGT GAATAGGCAGACAGACTTGTCACTGG | HPLC |
| TRBC-RT-BC012-1-P | /5phos/ctagcgtacgagcactccagattacttcca**ACTACTCA**ACCA GTGTGGCCTTTTGGGTGTGGGAG | HPLC |
| TRAC-RT-BC013-1-P | /5phos/ctagcgtacgagcactccagattacttcca**ATACGTCA**CGGT GAATAGGCAGACAGACTTGTCACTGG | HPLC |
| TRBC-RT-BC013-1-P | /5phos/ctagcgtacgagcactccagattacttcca**ATACGTCA**ACCA GTGTGGCCTTTTGGGTGTGGGAG | HPLC |
| TRAC-RT-BC014-1-P | /5phos/ctagcgtacgagcactccagattacttcca**ATCATGTG**CGGT GAATAGGCAGACAGACTTGTCACTGG | HPLC |
| TRBC-RT-BC014-1-P | /5phos/ctagcgtacgagcactccagattacttcca**ATCATGTG**ACCA GTGTGGCCTTTTGGGTGTGGGAG | HPLC |
| TRAC-RT-BC015-1-P | /5phos/ctagcgtacgagcactccagattacttcca**AACGCCGA**CGG TGAATAGGCAGACAGACTTGTCACTGG | HPLC |
| TRBC-RT-BC015-1-P | /5phos/ctagcgtacgagcactccagattacttcca**AACGCCGA**ACC AGTGTGGCCTTTTGGGTGTGGGAG | HPLC |
| TRAC-RT-BC016-1-P | /5phos/ctagcgtacgagcactccagattacttcca**ATTCGGTT**CGGT GAATAGGCAGACAGACTTGTCACTGG | HPLC |
| TRBC-RT-BC016-1-P | /5phos/ctagcgtacgagcactccagattacttcca**ATTCGGTT**ACCA GTGTGGCCTTTTGGGTGTGGGAG | HPLC |

**Supplemental Table 1.** Human TRAC and TRBC RT primer sequences. mRNA targeting regions shown in UPPERCASE, 10x Genomics sample barcode shown in **bold**, constant sequence and splint oligo overlapping sequence shown in lowercase.

| Name | Full Primer Sequence | Purification |
|---|---|---|
| Bdg-RT-BC001-38nt | CCTAAAGTTGGAAGTAATCTGGAGTGCTCGTA CGCTAGCGGTCCTAGCAA | HPLC |
| Bdg-RT-BC002-38nt | TTCCCGTTTGGAAGTAATCTGGAGTGCTCGTAC GCTAGCGGTCCTAGCAA | HPLC |
| Bdg-RT-BC003-38nt | AGCCTACTTGGAAGTAATCTGGAGTGCTCGTA CGCTAGCGGTCCTAGCAA | HPLC |
| Bdg-RT-BC004-38nt | GTCAACATTGGAAGTAATCTGGAGTGCTCGTA CGCTAGCGGTCCTAGCAA | HPLC |
| Bdg-RT-BC005-38nt | AGGTCTGTTGGAAGTAATCTGGAGTGCTCGTA CGCTAGCGGTCCTAGCAA | HPLC |
| Bdg-RT-BC006-38nt | GTTGGGATTGGAAGTAATCTGGAGTGCTCGTA CGCTAGCGGTCCTAGCAA | HPLC |
| Bdg-RT-BC007-38nt | CTCTACTTTGGAAGTAATCTGGAGTGCTCGTAC GCTAGCGGTCCTAGCAA | HPLC |
| Bdg-RT-BC008-38nt | TCACAGCTTGGAAGTAATCTGGAGTGCTCGTA CGCTAGCGGTCCTAGCAA | HPLC |
| Bdg-RT-BC009-38nt | CAGACTGTTGGAAGTAATCTGGAGTGCTCGTA CGCTAGCGGTCCTAGCAA | HPLC |
| Bdg-RT-BC010-38nt | CACTCACTTGGAAGTAATCTGGAGTGCTCGTA CGCTAGCGGTCCTAGCAA | HPLC |
| Bdg-RT-BC011-38nt | TTGCCTCTTGGAAGTAATCTGGAGTGCTCGTAC GCTAGCGGTCCTAGCAA | HPLC |
| Bdg-RT-BC012-38nt | TGAGTAGTTGGAAGTAATCTGGAGTGCTCGTA CGCTAGCGGTCCTAGCAA | HPLC |
| Bdg-RT-BC013-38nt | TGACGTATTGGAAGTAATCTGGAGTGCTCGTA CGCTAGCGGTCCTAGCAA | HPLC |
| Bdg-RT-BC014-38nt | CACATGATTGGAAGTAATCTGGAGTGCTCGTA CGCTAGCGGTCCTAGCAA | HPLC |
| Bdg-RT-BC015-38nt | TCGGCGTTTGGAAGTAATCTGGAGTGCTCGTA CGCTAGCGGTCCTAGCAA | HPLC |
| Bdg-RT-BC016-38nt | AACCGAATTGGAAGTAATCTGGAGTGCTCGTA CGCTAGCGGTCCTAGCAA | HPLC |

**Supplemental Table 2.** hTRAC and hTRBC bridging sequences used for mediating splinting and ligation of first strand cDNA onto 10X Genomics GEM Beads for scRNAseq capture

| Name | Variable domain binding primer region |
|---|---|
| TRAV1 | AGGTCGTTTTTCTTCATTCCTTAGTC |
| TRAV2 | ACGATACAACATGACCTATGAACGG |
| TRAV3.1 | CTTTGAAGCTGAATTTAACAAGAGCC |
| TRAV4.1 | CTCCCTGTTTATCCCTGCCGAC |
| TRAV5.1 | AAACAAGACCAAAGACTCACTGTTC |
| TRAV6 | AAGACTGAAGGTCACCTTTGATACC |
| TRAV7 | ACTAAATGCTACATTACTGAAGAATGG |
| TRAV8 | GCATCAACGGTTTTGAGGCTGAATTTAA |
| TRAV8.1 | GCATCAAGGGCTTTGAGGCTGAATTTAT |
| TRAV8.3 | GCATTAAAGGCTTTGAGGCTGAATTTAA |
| TRAV9 | GAAACCACTTCTTTCCACTTGGAGAA |
| TRAV10 | TACAGCAACTCTGGATGCAGACAC |
| TRAV12 | GAAGATGGAAGGTTTACAGCACA |
| TRAV13.1 | GACATTCGTTCAAATGTGGGCGAA |
| TRAV13.2 | GGCAAGGCCAAAGAGTCACCGT |
| TRAV14 | TCCAGAAGGCAAGAAAATCCGCCA |
| TRAV16 | GCTGACCTTAACAAAGGCGAGACA |
| TRAV17 | TTAAGAGTCACGCTTGACACTTCCA |
| TRAV18 | GCAGAGGTTTTCAGGCCAGTCCT |
| TRAV19 | TCCACCAGTTCCTTCAACTTCACC |
| TRAV20 | GCCACATTAACAAAGAAGGAAAGCT |
| TRAV21 | GCCTCGCTGGATAAATCATCAGGA |
| TRAV22 | ACGACTGTCGCTACGGAACGCTA |
| TRAV23 | CACAATCTCCTTCAATAAAAGTGCCA |
| TRAV24 | ACGAATAAGTGCCACTCTTAATACCA |
| TRAV25 | GTTTGGAGAAGCAAAAAAGAACAGCT |
| TRAV26.1 | CAGAAGACAGAAAGTCCAGCACCT |
| TRAV26.2 | ATCGCTGAAGACAGAAAGTCCAGT |
| TRAV27 | ACTAACCTTTCAGTTTGGTGATGCAA |
| TRAV29 | CTTAAACAAAAGTGCCAAGCACCTC |
| TRAV30 | AATATCTGCTTCATTTAATGAAAAAAAGC |
| TRAV34 | CCAAGTTGGATGAGAAAAAGCAGCA |
| TRAV35 | CTCAGTTTGGTATAACCAGAAAGGA |
| TRAV36 | GGAAGACTAAGTAGCATATTAGATAAG |

| | |
|---|---|
| TRAV38 | CTGTGAACTTCCAGAAAGCAGCCA |
| TRAV39 | CCTCACTTGATACCAAAGCCCGT |
| TRAV40 | AGGCGGAAATATTAAAGACAAAAACTC |
| TRAV41 | GATTAATTGCCACAATAAACATACAGG |
| TRBV2 | GCCTGATGGATCAAATTTCACTCTG |
| TRBV3-1 | TCTCACCTAAATCTCCAGACAAAGCT |
| TRBV4 | CCTGAATGCCCCAACAGCTCTC |
| TRBV5-1 | CGATTCTCAGGGCGCCAGTTCTCT |
| TRBV5-4,8 | CTCTGAGCTGAATGTGAACGCCT |
| TRBV6-1 | TGGCTACAATGTCTCCAGATTAAACAA |
| TRBV6-2,3 | CCCTGATGGCTACAATGTCTCCAGA |
| TRBV6-4 | GTGTCTCCAGAGCAAACACAGATGATT |
| TRBV6-5,6 | GTCTCCAGATCAACCACAGAGGAT |
| TRBV6-8 | GTCTCTAGATTAAACACAGAGGATTTC |
| TRBV6-9 | GGCTACAATGTATCCAGATCAAACA |
| TRBV7-2 | TCGCTTCTCTGCAGAGAGGACTGG |
| TRBV7-3 | CGGTTCTTTGCAGTCAGGCCTGA |
| TRBV7-4,6 | TCTCCACTCTGAAGATCCAGCGCA |
| TRBV7-4,6 | TCTCCACTCTGACGATCCAGCGCA |
| TRBV7-7 | GCAGAGAGGCCTGAGGGATCCAT |
| TRBV7-8 | CCAGTGATCGCTTCTTTGCAGAAA |
| TRBV7-9 | CTGCAGAGAGGCCTAAGGGATCT |
| TRBV9 | CTCCGCACAACAGTTCCCTGACTT |
| TRBV10-1,3 | CAGATGGCTACAGTGTCTCTAGATCAAA |
| TRBV10-1,3 | CAGATGGCTATAGTGTCTCTAGATCAAA |
| TRBV10-2 | GTTGTCTCCAGATCCAAGACAGAGAA |
| TRBV11 | GCAGAGAGGCTCAAAGGAGTAGACT |
| TRBV12-3,4 | GCTAAGATGCCTAATGCATCATTCTC |
| TRBV12-5 | CTCAGCAGAGATGCCTGATGCAACT |
| TRBV13 | TCTCAGCTCAACAGTTCAGTGACTA |
| TRBV14 | GCTGAAAGGACTGGAGGGACGTAT |
| TRBV15 | GATAACTTCCAATCCAGGAGGCCG |
| TRBV16 | GCTAAGTGCCTCCCAAATTCACCC |
| TRBV18 | GGAACGATTTTCTGCTGAATTTCCCA |
| TRBV19 | GGTACAGCGTCTCTCGGGAGAAGA |

| | |
|---|---|
| TRBV20-1 | GGACAAGTTTCTCATCAACCATGCAA |
| TRBV24-1 | TGGATACAGTGTCTCTCGACAGGC |
| TRBV25-1 | CAACAGTCTCCAGAATAAGGACGGA |
| TRBV27-1 | TACAAAGTCTCTCGAAAAGAGAAGAGGA |
| TRBV28 | GGGGTACAGTGTCTCTAGAGAGA |
| TRBV29 | GTTTCCCATCAGCCGCCCAAACCTA |
| TRBV30 | CAGACCCCAGGACCGGCAGTTCAT |
| | Full variable primer example (inclusive of partial TruSeq2 overhang, shown in lowercase) |
| TRAV38 | cagacgtgtgctcttccgatctCTGTGAACTTCCAGAAAGCAGCCA |
| TRBV14 | cagacgtgtgctcttccgatctGCTGAAAGGACTGGAGGGACGTAT |

**Supplemental Table 3.** hTRAV sequences and example full length primers used during the pre-amplification stage of library prep for enriching full length TCRalpha and beta CDR3.

| **Donor** | **Allele 1** | **Allele 2** | **Allele 1** | **Allele 2** |
|---|---|---|---|---|
| **Donor 1** | HLA-A*02:01 | HLA-A*24:02 | HLA-DRB1*15:01 | HLA-DRB1*15:02 |
| **Donor 2** | HLA-A*02:01 | HLA-A*11:02 | HLA-DRB1*04:03:01 | HLA-DRB1*15:02 |
| **Donor 3** | HLA-A*01:01 | HLA-A*02:03 | DRB1*07:01 | DRB1*08:03 |

**Supplemental Table 4.** Donor HLA typing

| id | name | read | pattern | sequence | feature_type |
|---|---|---|---|---|---|
| ADT_C0053 | Hu.CD11c | R2 | 5PNNNNNNNNNN(BC) | TACGCCTATAACTTG | Antibody Capture |
| ADT_C0081 | Hu.CD14_M5E2 | R2 | 5PNNNNNNNNNN(BC) | TCTCAGACCTCCGTA | Antibody Capture |
| ADT_C0083 | Hu.CD16 | R2 | 5PNNNNNNNNNN(BC) | AAGTTCACTCTTTGC | Antibody Capture |
| ADT_C0050 | Hu.CD19 | R2 | 5PNNNNNNNNNN(BC) | CTGGGCAATTACTCG | Antibody Capture |
| ADT_C0034 | Hu.CD3_UCHT1 | R2 | 5PNNNNNNNNNN(BC) | CTCATTGTAACTCCT | Antibody Capture |
| ADT_C0048 | Hu.CD45_2D1 | R2 | 5PNNNNNNNNNN(BC) | TCCCTTGCGATTTAC | Antibody Capture |
| ADT_C0072 | Hu.CD4_RPA.T4 | R2 | 5PNNNNNNNNNN(BC) | TGTTCCCGCTCAACT | Antibody Capture |
| ADT_C0047 | Hu.CD56 | R2 | 5PNNNNNNNNNN(BC) | TCCTTTCCTGATAGG | Antibody Capture |
| ADT_C0046 | Hu.CD8 | R2 | 5PNNNNNNNNNN(BC) | GCGCAACTTGATGAT | Antibody Capture |
| DEX_01 | HLA-A*0201+NLVPMVATV+pp65+CMV | R2 | 5PNNNNNNNNNN(BC) | GCCGCGCGGTAGCGC | Antibody Capture |

**Supplemental Table 5.** Antibody Derived Tag CellRanger Multi Table, including ADT probe name, protein target, read, tag pattern, sequence, and feature type.

| Target | Conjugate | Clone | Purpose |
|---|---|---|---|
| CD3 | BUV395 | UCHT1 | Pan T cells |
| CD56 | BUV615 | NCAM16.2 | NK cell Exclusion |
| CD19 | BUV615 | HIB19 | B Cell Exclusion |
| CD8 | BUV737 | RPA-T8 | CD8 T cells |
| pp65+ | PE | N/A | Dextramer |
| Live/Dead | FVS510 | N/A | Viability |
| CD4 | BB700 | SK3 | CD4 T cells |
| CD14 | SparkYG 593 | S18004B | Monocyte Exclusion |

**Supplemental Table 6.** Antibody panel used in dextramer flow analysis
